# Nanocrystalline silver layer of knitted polyester outperforms other silver-containing wound dressings in an in vitro wound model

**DOI:** 10.1101/2023.01.23.524940

**Authors:** Jonathan Hus, Ricardo F. Frausto, Joel Grunhut, Nir J. Hus

**Author notes:** Correspondence: Nir Hus MD, Ph.D., FACS., Assistant Professor of Surgery, Schmidt College of Medicine, Florida Atlantic University, FL, 4600 Linton Blvd suit #100, Delray Beach, FL 33445.

## Abstract

**Background:** Silver possesses cytotoxic properties against many microorganisms and is regularly used in wound care. Current evidence supporting the use of one type of silver-containing wound dressing (SCWD) is insufficient.

**Methods:** To examine the ability of selected SCWDs to inhibit the growth of two strains of bacteria (*E. coli and S. aureus*) commonly found in wounds, an in vitro wound model was used. Bacteria were applied to the surface of nutrient agar and a piece of each SCWD was applied to the bacteria. The plates were incubated at 37°C overnight. The zone of inhibition (ZI) around each SCWD was measured in cm^2^.

**Results:** The mean ZI for Acticoat Flex-3 on *E. coli* was 1.59 ± 0.15 cm^2^, which was significantly greater than that observed for Aquacel Ag (p<0.001), Mepilex Ag (p<0.0001), Mepitel Ag (p<0.001), Optifoam (p<0.0001) and Tegaderm Alginate Ag (p<0.01), but statistically indistinguishable from Maxorb II Ag. The mean ZI on *S.aureus* was 1.21 ± 0.16 cm^2^, which was greater than Aquacel Ag (p<0.05), Mepilex (p<0.0001), Optifoam (p<0.0001) and Tegaderm Alginate Ag (p<0.05), but statistically indistinguishable from Maxorb II Ag or Mepitel Ag.

**Conclusions:** Of the SCWDs tested, Acticoat Flex-3 demonstrated the most robust antimicrobial effect. Herein we show that Acticoat Flex-3 may provide the most wound protection against bacterial infection. In conclusion, these data provide clinicians with additional independent evidence to inform their clinical practice on the use of specific wound dressings.

**PRECIS:** The antimicrobial properties of Acticoat Flex-3, a dressing composed of nanocrystalline silver layer of knitted polyester, outperformed other silver-containing dressings against *E. coli* and *S. aureus*.

## INTRODUCTION

The utilization of silver in wound care has been reported as early as 69 B.C.^1^ Silver’s properties exert cytotoxic effects on fungal, bacterial, and viral microbes.^2,3^ The spectrum of bacterial coverage is wide and includes *Staphylococcus aureus* and *Escherichia coli*, two of the most commonly implicated bacteria in wound infections.^4^ Over the last two decades, studies on wound care products containing silver suggest silver fulfills a valuable role in wound care.^5^ Silver-containing wound dressings (SCWD) were designed to help decrease wound infection and have transformed the scope of wound care.

Silver must be in a soluble form for it to be biologically active.^6^ Silver ions (Ag^+^) provides cytotoxic activity through the interruption of biofilms, the increased uptake of antibiotics, and the generation of reactive oxygen species.^7–10^ Earlier SCWDs provided a quick initial load of Ag^+^ and would quickly become depleted with prolonged contact with chloride ions in serum. New SCWDs have been designed to release a slow and steady supply of Ag^+^. For example, dressings under the Acticoat brand contain Ag^0^ clusters. These silver clusters were designed to slowly release silver upon contact with wound fluid.^6^

Previous studies have shown differences in wound treatment outcomes between a variety of wound dressings such as a silver hydrogel dressing, PolyMem Silver, and Acticoat.^2^ An additional study found differences in antimicrobial activity amongst Acticoat, Acticoat Moisture Control, Acticoat Absorbent, SilvercelTM, Aquacel Ag, Urgotul SSD, and Actisorb.^11^ The authors reason that the differences were the result of disproportionality in silver concentrations since higher concentrations correlated with more robust antibacterial properties.^11,12^

Acticoat Flex-3, an Ag^0^ silver nanoparticle-based dressing, has been shown to be an effective antimicrobial dressing and has demonstrated effective wound healing properties.^13,14^In a previous study comparing a limited number of other SCWDs, Acticoat Flex-3 had a greater silver release over Actisorb Silver 220, Aquacel Ag, and Mepilex Ag.^14^ However, the authors did not investigate their respective antimicrobial properties.^15^ Although SCWDs with higher silver release, not to be confused with higher starting concentration, should conceptually exhibit higher antimicrobial effects, yet this has not been previously shown.

Among newer SCWDs, there is limited evidence that one SCWD significantly outperforms others. The different SCWDs each contain unique materials, varied levels of silver, and unique compositions of silver. This can potentially result in different antimicrobial capacities. With continued concerns about multidrug-resistant bacteria as a consequence of prescribing antibiotics to fight infections, a renewed interest in optimizing silver–based wound therapies has emerged.^16^

In the absence of clear experimental evidence, a combination of factors dictate the clinical use of a particular dressing. These include the availability of the dressing, the familiarity of the physician with the dressing, and the type of wound. For patients to achieve better wound care outcomes, independent studies assessing the effectiveness of different SCWDs are needed. Herein, using an in vitro wound model, we demonstrate that the antibacterial properties of Acticoat Flex-3 outperform those of other industry-leading SCWDs.

## METHODS AND MATERIALS

### Cell density measurements

Colonies of *E. coli (ATCC25922 Seattle 1946)* and *S. aureus (HIP10787 mupA positive QC strain Methycilin Resistant)* sourced from Thermo Scientific (LENEXA, KS 66215 USA) were independently inoculated into a 10 ml Luria Broth (LB) medium (Aldon Corporation Rochester, NY). These were grown overnight to saturation in a 37°C incubator and shaken vigorously at 250 cycles per minute on a rotary shaker. 10 μl of each saturated broth were then used to independently inoculate another 10 ml of an LB medium of *E. coli* and *S. aureus*. Thereafter, the broth cultures were grown until an optical density (OD) setting of 0.595 nm reached 0.6 as measured using a spectrophotometer (CGOLDENWALL 722N Visible Spectrophotometer) in accordance with National Committee for Clinical Laboratory Standards recommendations.

### Preparation of in vitro wound model (modified Kirby-Bauer Test)

Agar plates containing bacterial lawns were prepared by spreading 100 μL of 1:1000 diluted *E.coli* or *S. aureus*. Pieces of each SCWD measuring 1 cm^2^ were cut using sterile industrial-grade scissors. Subsequently, the pieces of SCWD were placed on the LB agar plates onto which bacteria had been spread. Sterile gauze was used as a negative control. Whatman paper (3M Saint Paul, MN) cut to 1 cm^2^ and pre-wetted with 100 μg/mL gentamicin was used as a positive control. The plates were incubated overnight at 37°C then images were acquired. The area of the zones of inhibition (ZI) was measured for each dressing and plotted as cm^2^.

### Unbiased Semiquantitative Analysis

As a summary of the quantitative results obtained using the modified Kirby-Bauer Test, we performed a semiquantitative analysis. Each data point corresponding to one of four combinations of bacterial strain and zone (*E. coli*/ZI, and S. aureus/ZI) was plotted and the quartiles were determined for each combination (Supplemental Data Fig. 1). To avoid skewness in the data, zeroes were removed from this analysis. The mean ZI for each SCWD was assigned a number of plus signs (+) based on the quartile range in which it was found. The first/bottom quartile (0-25%) was given a single plus sign, the second quartile (25-50%) was given two plus signs, the third quartile (50-75%) was given three plus signs, and the fourth/top quartile (75-100%) was given four plus signs. For scoring purposes, the plus sign was defined as one point.

### Statistical Analysis

The ZI of each SCWD was measured using ImageJ software from the National Institutes of Health. An image of each plate in the same position was captured and measured in ImageJ by tracing the clear zones (inhibition or ZI) of each SCWD. The calculated values for the areas comprising the respective zones were subsequently graphed using GraphPad Prism 9 (GraphPad Software, Inc.). The values were statistically analyzed using one-way ANOVA and post hoc Tukey test for multiple comparisons. Statistically significant differences were defined as p<0.05.

## RESULTS

To examine the inhibitory effect of the SWCDs on *E. coli* and *S. aureus*, we performed a modified Kirby-Bauer Test. We obtained representative images for each of the SCWDs for the two strains of bacteria tested (Fig. 1). With respect to the effect of the SCWDs on *E. coli* growth, Acticoat Flex 3 showed the most robust inhibitory effect with a mean ZI of 1.59 ± 0.15 cm^2^ (Fig. 2A). The mean ZI of Aquacel Ag (0.89 ± 0.15 cm^2^) was significantly lower than that of Acticoat Flex 3 (p<0.01)(Fig. 2A). The mean ZI of Maxorb II Ag (1.43 ± 0.09 cm^2^) was not statistically significantly lower (p=0.96) compared with Acticoat Flex 3. Mepilex Ag (0.08 ± 0.01 cm^2^) had a significantly lower mean ZI than that of Acticoat Flex 3 (p<0.01). Similarly, Mepitel Ag (0.90 ± 0.03 cm^2^, p<0.001), Optifoam (0.37 ± 0.03 cm^2^, p<0.0001), Tegaderm Alginate Ag (0.96 ± 0.10 cm^2^, p<0.01) each had a mean ZI significantly lower than that of Acticoat Flex 3.

**Figure 1.**
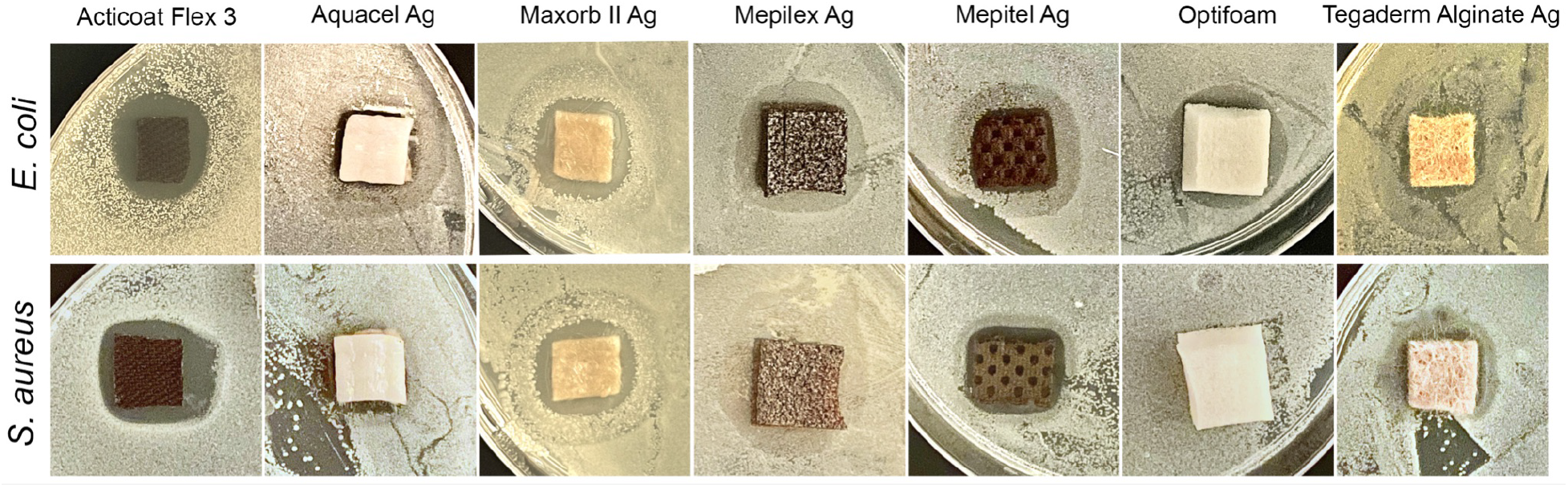
SCWDs demonstrate variable antimicrobial effects against *E. coli* and *S. aureus*. Representative images of each SCWD for *E. coli* and *S. aureus*. The zone of inhibition is defined by clear zones adjacent to the SCWD.

**Figure 2.**
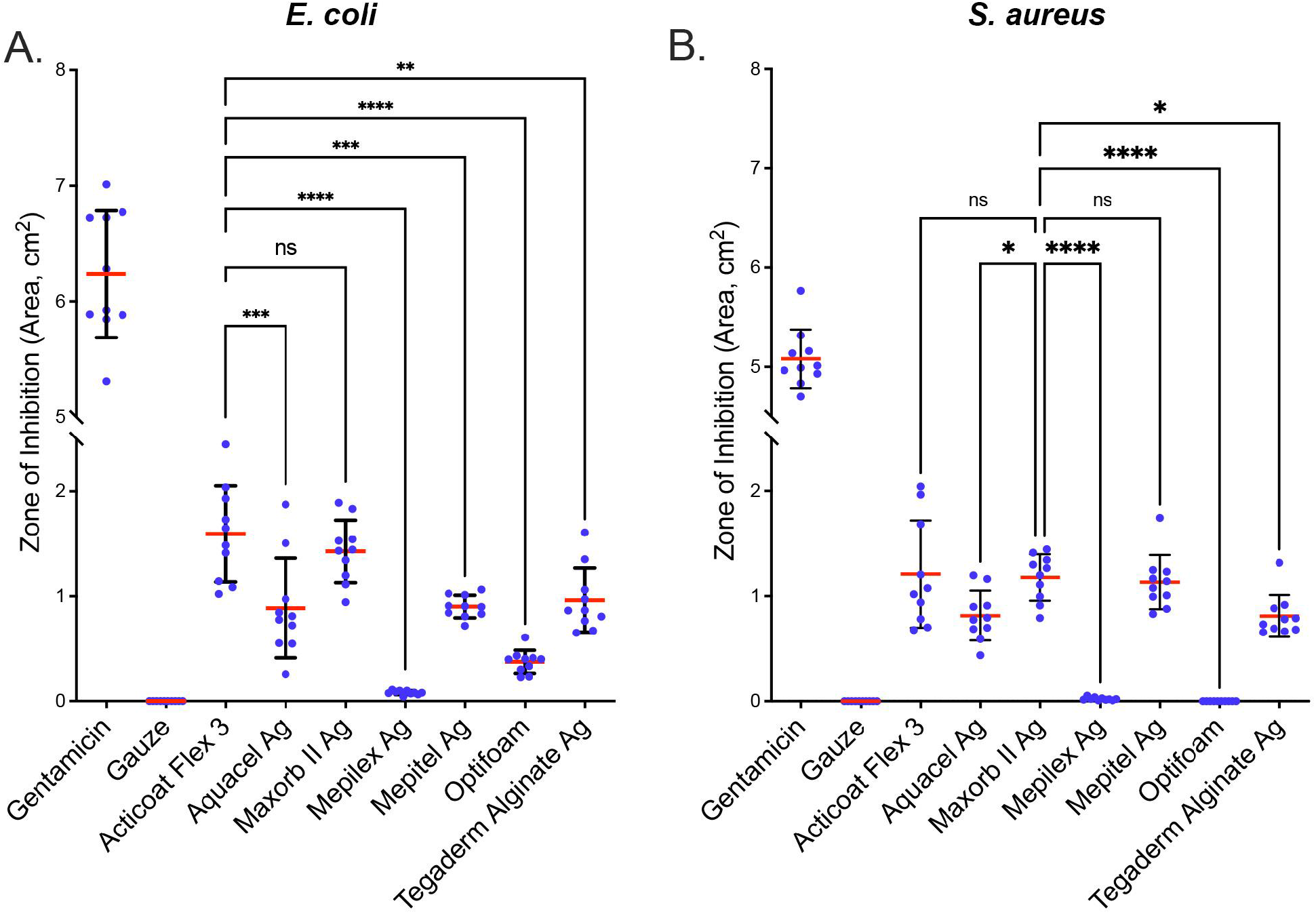
SCWDs differ in their antibacterial effect. Antibacterial effect of SCWDs against (A) *E. coli* and (B) *S. aureus*. Gentamicin was used as a positive control and gauze only was used as a negative control. Scatter dot plot with zone of inhibition given in cm^2^. Ten replicates (n=10) were performed for each SCWD. Statistical analysis was performed using ordinary one-way ANOVA with Tukey’s multiple comparisons test. Error bars = mean±SD. Mean is represented as a red line. Not significant, ns; **p<0.01, ***p<0.001 and ****p<0.0001.

The effect of the SCWDs on the growth of *S. aureus*, was also examined (Fig. 2B). We obtained the highest mean ZI for Maxorb II Ag, therefore the results and statistical analyses described here were obtained using Maxorb II Ag as the reference. The mean ZI for Acticoat Flex 3 (1.21 ± 0.16 cm^2^) was statistically indistinguishable from that obtained with Maxorb II Ag (1.18 ± 0.07 cm^2^, p=0.99). The mean ZI for Aquacel Ag (0.82 ± .08 cm^2^, p<0.05), Mepilex Ag (0.02 ± 0.01 cm^2^, p<0.0001), Optifoam ((0.00 ± 0.00 cm^2^, p<0.0001) and Tegaderm Alginate Ag (0.81 ± 0.06 cm^2^ p<0.05) were significantly lower than Maxorb II Ag (1.18 ± 0.07 cm^2^)). The mean ZI for Mepitel Ag (1.13 ± 0.08 cm^2^) was statistically indistinguishable from the mean ZI for Maxorb II Ag (1.18 ± 0.07 cm^2^, p=0.99).

For a comprehensive comparison of the results described above, we performed a semiquantitative analysis (Table 2). Acticoat Flex-3 had a score of 8 points (1 point for each “+”, maximum of 4 points per condition) with the maximum of 4 points for each condition. Maxorb II Ag had a score of 7 each and also showed a fairly strong antibacterial effect against both *E. coli* and *S. aureus*. Mepitel Ag (6 points), Aquacel Ag (5 points) and Tegaderm Alginate Ag (5 points) overall moderate antibacterial effects. Strikingly, Mepilex Ag (2 points) and Optifoam (1 point) showed the weakest antibacterial effects, with the former showing 1 point for each bacterial strain and the latter showing 1 and zero points for *E. coli* and *S. aureus*, respectively.

**Table 1.** Characteristics of silver-containing wound dressings.

| Product | Manufacturer | Composition | properties | Product number |
| --- | --- | --- | --- | --- |
| Acticoat Flex 3 | Smith & Nephew,<br>London, UK | Silver layer of Knitted polyester<br>Contains .69 mg/cm<sup>2</sup> - 1.64 mg/cm<sup>2</sup><br>Silver nanocrystalline structure (Ag<sup>0</sup>, slow release, converts to Ag<sup>+</sup>) | Will adhere to wound bed and aid in minor debreameant with the removal of the dressing. | 66800417 |
| Aquacel Ag | ConvaTec,<br>Greensboro NC | Silver-coated Hydrofiber sodium carboxymethylcellulose and 1.2% Ag<sup>+</sup> | Absorbs exudate and forms a tight seal surrounding the wound. | 412010 |
| Maxorb II Ag | Manufactured in China for Medline Industries Inc. Northfield IL. | Highly Absorbent Silver-coated foam pad<br>Contains a maximum of 0.306 mg of ionic silver per square cm, silver sodium hydrogen zirconium phosphate, and 100% calcium Alginate (Ag<sup>+</sup>) | Manages exudate and bacterial burden | MSC9945EP |
| Mepilex Ag | Mölnlycke Health Care AB, Göteborg Sweden | Silver-coated foam with silicone interface | Adheres primarily to the skin surrounding the wound and not to the wound bed itself | 287500 |
| Mepitel Ag | Mölnlycke Health Care AB, Göteborg Sweden | Silver-coated silicone<br>Contains 0.13 % by weight of Silver Chloride (Ag<sup>+</sup>) | Adheres to the skin and not to the wound bed | 391090 |
| Optifoam Ag Non-Adhesive | Manufactured in UK for Medline Industries Inc. Mundelein, IL. | Non-staining Antimicrobial Silver-coated pad with silicone adhesive border | Manages bacterial burden and absorbs exudate | MSC9614EP |
| 3M Tegaderm Alginate Ag | 3M inc. Saint Paul Minnesota | Silver-coated fiber<br>Contains silver sodium hydrogen zirconium phosphate (Ag<sup>+</sup>) | Absorbs exudate | 90303 |

**Table 2.** Unbiased semiquantitative analysis of bacterial inhibition.

| SCWD | *E. coli* | *S. aureus* | Score |
| --- | --- | --- | --- |
| Acticoat Flex 3 | ++++ | ++++ | 8 |
| Aquacel Ag | +++ | ++ | 5 |
| Maxorb II Ag | ++++ | +++ | 7 |
| Mepilex Ag | + | + | 2 |
| Mepitel Ag | +++ | +++ | 6 |
| Optifoam | + | - | 1 |
| Tegaderm Alginate Ag | +++ | ++ | 5 |

## DISCUSSION

Bacterial infection is a common occurrence that complicates wound healing. In the United States and Europe,the world’s largest wound-dressing markets, there has been an increasingly high demand for wound healing products where in 2014, the global annual cost for wound care was averaged at around $2.8 billion.^17^ Additionally, Medicare cost estimates for chronic and acute wound treatments in 2018 were between $28.1 billion to $96.8 billion.^17^ Silver-containing wound dressings (SCWDs) are an attractive and practical choice for wound treatment as silver has been shown to possess antimicrobial properties. Silver is detrimental to bacteria in part through its ability to damage the bacterial cell wall (resulting in increased membrane permeability), block enzyme and solute transport systems, prevent DNA and RNA replication, and block cellular respiration.^18,19^ During wound healing, microorganisms such as *S. aureus* and *E. coli* can predominate.^20^ Our comparative study demonstrated the variability in antimicrobial effectiveness of selected SCWDs against these bacteria. Nevertheless, Acticoat Flex-3 demonstrated the most consistent antimicrobial properties. Other studies have shown the effectiveness of different Acticoat-branded dressings, however, the relative effectiveness of Acticoat Flex-3 in an in vitro wound model in an in vitro wound model, in particular, has not been investigated.^11^

In 2007, Castellano et al. demonstrated that several Acticoat-branded wound dressings (also containing Ag), in addition to Aquacel Ag, were effective at inhibiting the growth of several strains of bacteria (*Staphylococcus aureus, Streptococcus faecalis, Escherichia coli, and Pseudomonas aeruginosa*).^11^ Similar to our study, Aquacel Ag performed at a moderate level compared with the other dressings tested. In our study, the most robust inhibitory effect against *S. aureus* was observed with Acticoat Flex-3, Maxorb II Ag, and Mepitel Ag, which were statistically indistinguishable from each other. Overall, the inhibitory effects of the selected SCWDs show that a majority of silver-containing wound dressings can negatively impact bacterial growth. Taken together, we show through our comparative study that the antimicrobial effectiveness of SCWDs commonly used in current clinical settings varies significantly. Studies show that Acticoat-branded SCWDs (e.g., Acticoat Flex 3) provide a slower release and prolonged exposure to bioactive silver cations as a result of their proprietary nanocrystalline composition.^21^ Because of this variability, we performed a semi-quantitative analysis that identified Acticoat Flex 3 as the most consistent antimicrobial SCWD in our study. The semi-quantitative analysis showed that Acticoat Flex 3 demonstrated the most robust antimicrobial effectiveness across conditions, but Maxorb II Ag was also effective across conditions to a marginally lesser extent. Aquacel Ag, Mepitel Ag and Tegaderm Alginate Ag performed moderately well, while Mepilex Ag and Optifoam performed poorly with a weak or no antimicrobial response against *E. coli* and *S. aureus*.

Many factors may explain the variability we observed with the selected SCWDs, but the factor that is likely to drive the biggest effect is the bioavailability of the Ag^+^ ions. In 2012, Rigo et al. reported the amounts and rate of silver released from a number of SCWDs that we also tested in our study (Acticoat Flex 3, Aquacel Ag, and Mepilex Ag).^14^ Initially, the authors independently verified the silver concentrations in the dressings and found that they were consistent with those provided by the vendors (1.379 ± 0.091 mg/cm^2^ for Acticoat Flex 3, 0.993 ± 0.078 mg/cm^2^ for Mepilex Ag, and 0.111 ± 0.004 mg/cm^2^ for Aquacel Ag). Interestingly, these concentrations do not correlate with the antimicrobial effects we observed. This suggests that other factors influence the effectiveness of a particular SCWD. The authors also measured the rate and amount of silver released by placing the dressings in several solutions, including a bioengineered serum substitute.^14^ They showed Acticoat Flex-3 and Aquacel Ag continued to release silver up to at least 3 days in the serum substitute. This was in contrast with Mepilex Ag, which has a lower concentration of Ag than Acticoat Flex 3, but nonetheless showed a very high initial rate of release and reached maximum concentration (greater than that achieved by Acticoat Flex 3 by day 3) within an hour. In our study, if we assume slow release from Acticoat Flex 3 and Aquacel Ag, then inhibition of *E. coli* and *S. aureus* inversely correlates with the rate of release but not with the amount of Ag in solution (or agar matrix in our case). Taken together, these data suggest that the degree to which Ag is bioavailable, which defines the antimicrobial profile of any SCWD, is subject to a combination of technology- and wound-specific variables.

In addition to in vitro studies, in vivo models have also been used to study the effectiveness of SCWDs. The advantage of these models is that they establish a more physiologically relevant wound environment. Recently, a study showed that SCWDs containing nanocrystalline silver (e.g., Acticoat Flex3) were superior to silver-plated dressings and non-SCWDs in an in vivo wound model.^22^ However these studies were limited in the number of products they compared. A review of the literature spanning the last ten years reveals a paucity of in vitro studies investigating the antibacterial efficacy of SCWDs.^15,20,23,24^ Additionally, earlier in vitro studies showed conflicting results.^24–26^ As such, our study represents the most comprehensive list of SCWDs investigated for their effectiveness at inhibiting bacterial growth.

## CONCLUSIONS

In considering the conclusions of our study we recognize a few study limitations. One such limitation is that our in vitro model does not fully recapitulate wounds in humans. However, this affords us the ability to study the effectiveness of the selected SCWDs in a straightforward and highly controlled environment. In addition, our study is limited by having examined only two bacterial strains, although wounds can contain a wide spectrum of bacteria, such as *Pseudomonas aeruginosa, Proteus mirabilis*, and *Streptococcus pyogenes*.^20^ Finally, one must also be aware of the potential cytotoxic effects that silver may have on keratinocytes and fibroblasts.^27^ Nevertheless, herein we show that Acticoat Flex 3 possessed the highest antibacterial properties compared with other contemporary SCWDs. However, Maxorb II Ag also showed robust antibacterial effects. In addition, our study provides valuable insight into the effectiveness of commonly used SCWDs that can be used as one factor to inform their clinical application.

---

**Figure S1.**
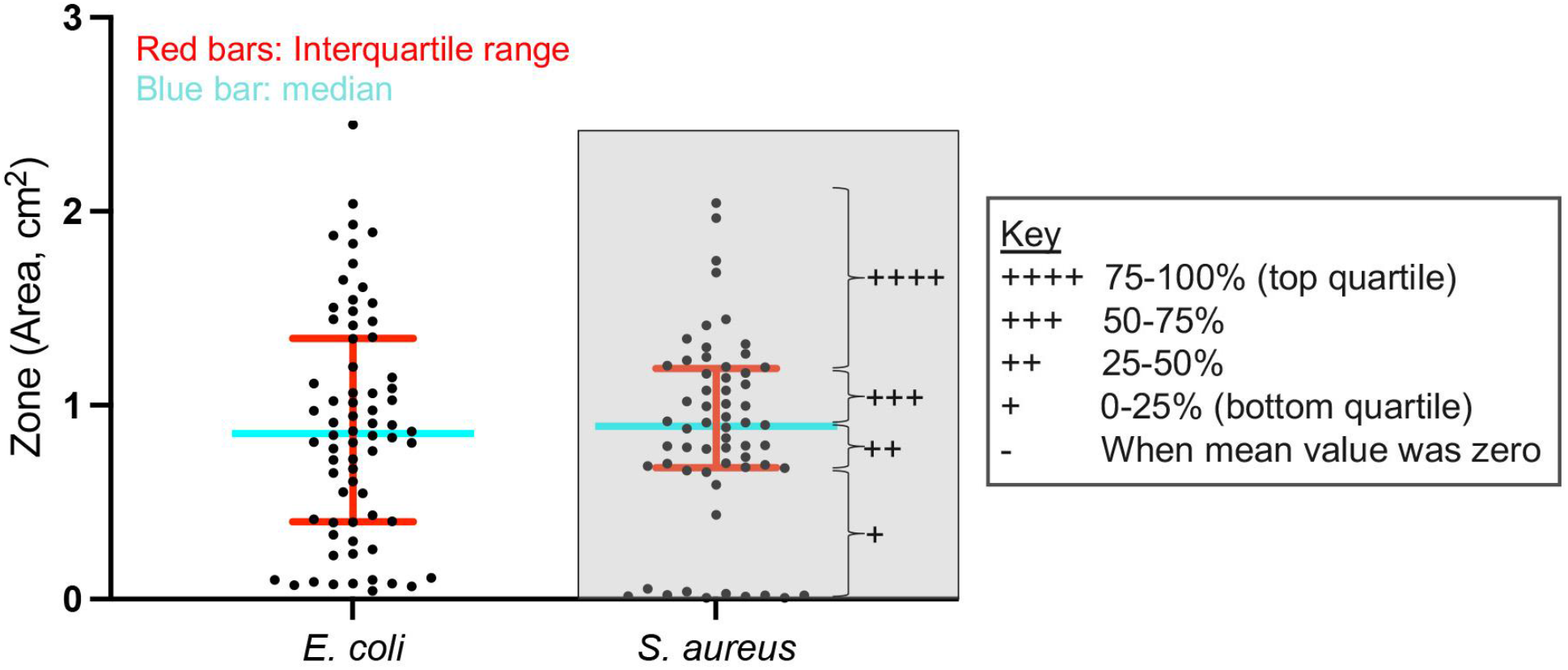
Defining the quartile ranges for semiquantitative analysis. To perform the unbiased semiquantitative analysis we first defined quartile ranges for all data grouped by bacterial strain. Scatter dot plot with all data from *E. coli* or *S. aureus*. To remove skewness in the data, zero values were removed from this analysis. Red bar represents the interquartile range comprising the center 50% of data. Blue bar represents the median, which separates the top 50% from the bottom 50% of data. The key shows the number of plus signs (“+”) assigned to each quartile. This is given as an example in the gray-filled box for *S. aureus*.

## AUTHOR CONTRIBUTIONS

J.H. and N.J.H. conceptualized the study. J.H drafted the manuscript and performed experiments. J.H. and R.F.F. designed experiments, interpreted results, analyzed data, and prepared figures/tables. J.H., R.F.F. and J.G. performed extensive review and critical revision of the manuscript. N.J.H provided reagents, consumables, and instruments. All authors reviewed and approved the manuscript.

